## Supplementary Figures for "Analyses of cell wall synthesis in *Clostridioides difficile* reveal a diversification in cell division mechanisms in endospore-forming bacteria"

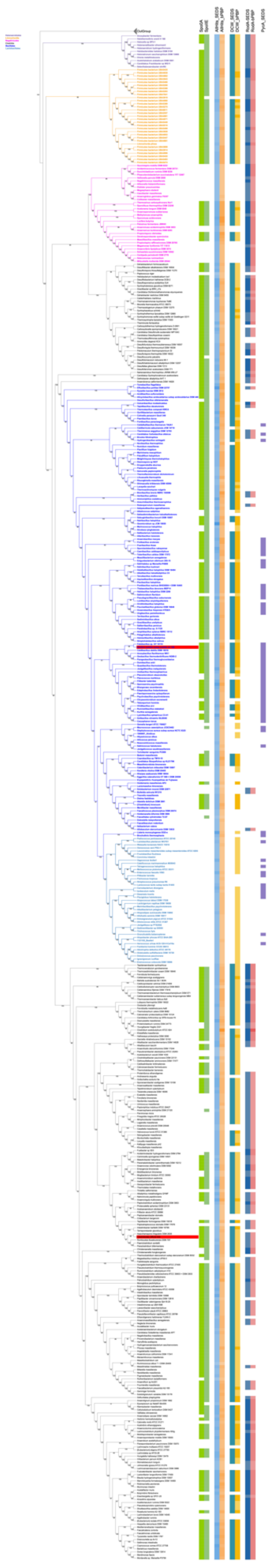

**Supplementary Fig. 1: Reference phylogeny of the Firmicutes with the mapping of the sporulation, SEDS glycosyltransferases, and bPBP homologs.** Maximum likelihood tree of the Firmicutes based on a supermatrix containing 497 taxa, 3,776 amino acid characters. The tree was inferred with IQ-TREE 1.6.3 using the LG+I+G4 model. Values on the nodes correspond to ultrafast bootstraps, computed on 1000 replicates. Halanaerobiales, Limnochordia, Negativicutes, Bacilli, and Clostridia are colored purple, orange, pink, blue, and black, respectively, in the tree. In red are highlighted *B subtilis* and *C difficile*. On the tree are mapped, respectively: the presence of *spo0A* and *spoIIE*; the presence of all SEDS glycosyltransferases and bPBP homologs found in the genome; the presence of *ftsW*/*spoVE* and *ftsI*/*spoVD* found in the *dcw* cluster; the presence of *rodA* and *mrdA* in pair; and the presence of the *ftsW* and *pycA* in synteny in the genome.

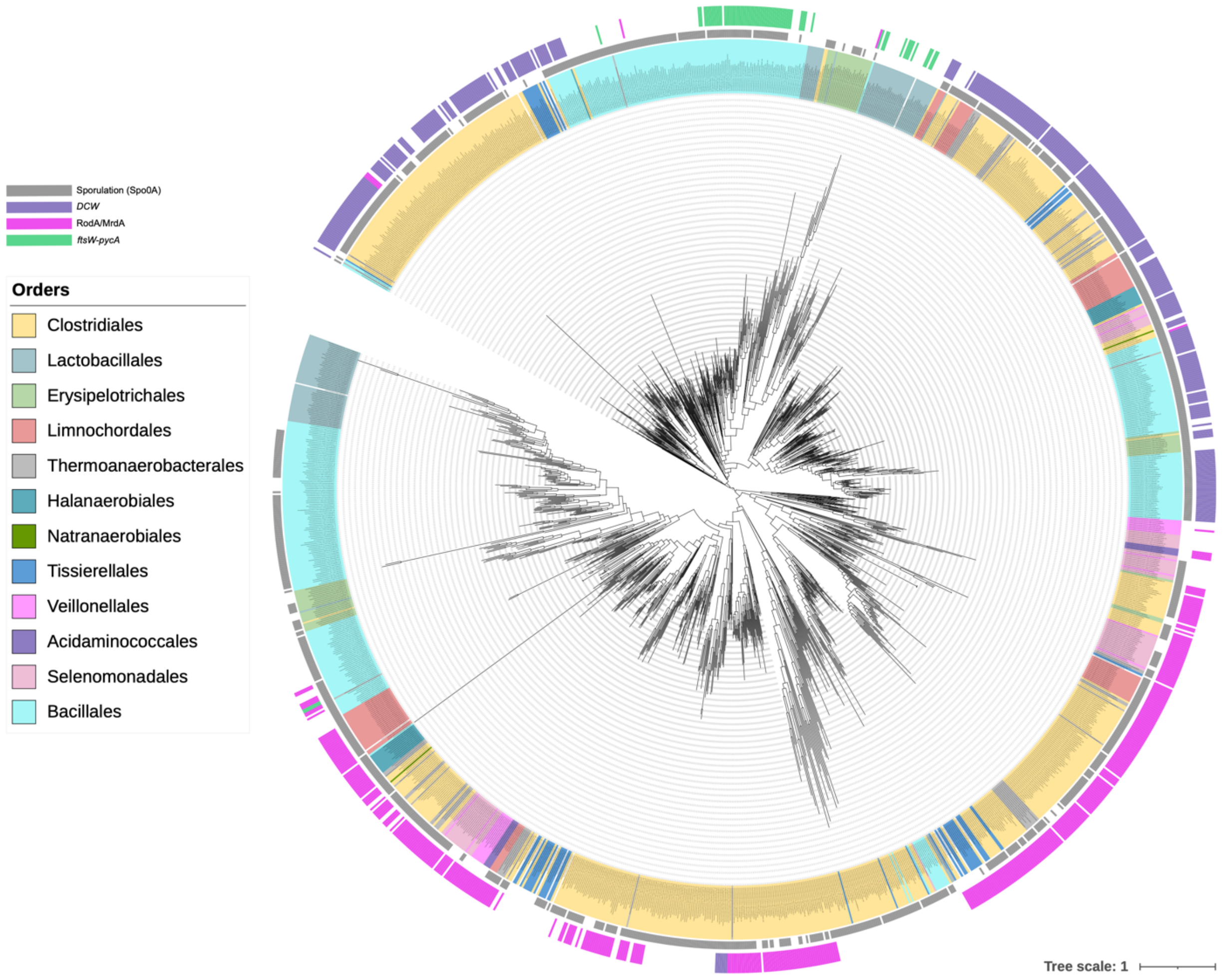


**Supplementary Fig. 2. Phylogenetic tree of SEDS homologs in the Firmicutes.** Maximum likelihood tree of all SEDS homologs based on an alignment containing 1,446 leaves and 343 amino acid characters. The tree was inferred with IQ-TREE2 using the LG+R10+I model according to the BIC criterium. On the tree are mapped the presence of *spo0A* (in red), and the synteny of SEDS hits in the *dcw* cluster (purple), paired with *mrdA* (pink), or paired with *pycA* (green). Scale bar corresponds to the average number of substitutions per site.

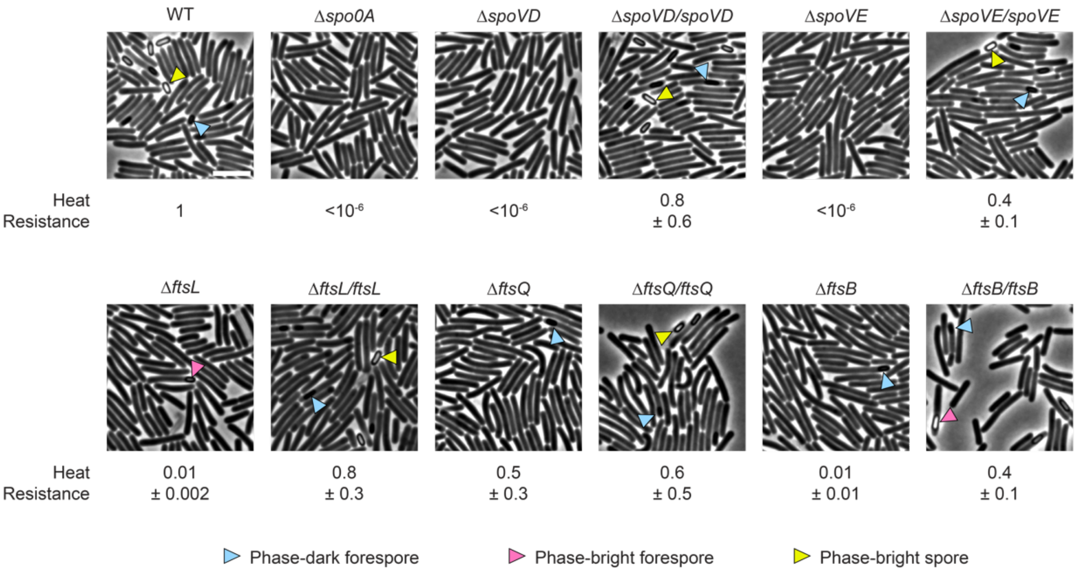


**Supplementary Fig. 3 | Sporulation efficiencies of mutant and complemented strains.** Representative phase-contrast micrographs of mutant and complemented cells isolated from sporulation-inducing media after 20-22 hours of growth. Immature phase-dark forespores are indicated by blue arrows, and mature phase-bright forespores and free spores are indicated by pink and yellow arrows, respectively. Scale bar, 5 µm. The mean efficiency of heat-resistant spore formation (heat resistance) relative to WT ± standard deviation is indicated below each strain. The Δ*spo0A* strain was used as a negative control because it does not initiate sporulation. Data from three independent experiments.

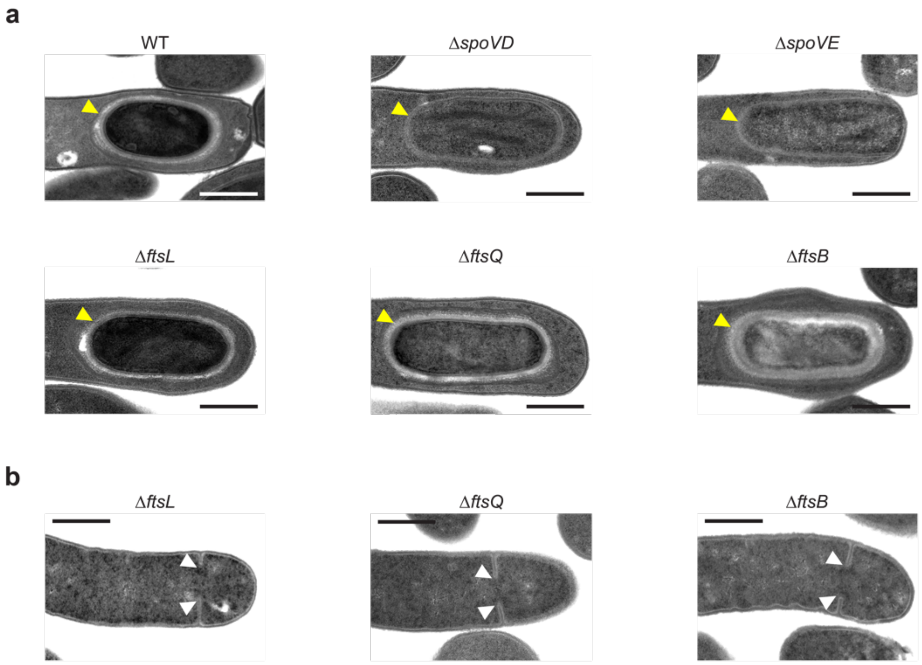


**Supplementary Fig. 4 | TEM analysis of sporulating cells. a**, Representative TEM micrographs showing engulfed forespores in WT, Δ*spoVD*, Δ*spoVE*, Δ*ftsL*, Δ*ftsQ*, and Δ*ftsB*. The presence (in WT, Δ*ftsL*, Δ*ftsQ*, and Δ*ftsB*) or absence (in Δ*spoVD*, and Δ*spoVE*) of a cortex layer is indicated (yellow arrows). Scale bars, 500 nm. **b**, Representative TEM micrographs of Δ*ftsL*, Δ*ftsQ*, and Δ*ftsB* cells that initiate but fail to complete septum formation (white arrows). Scale bars, 500 nm.

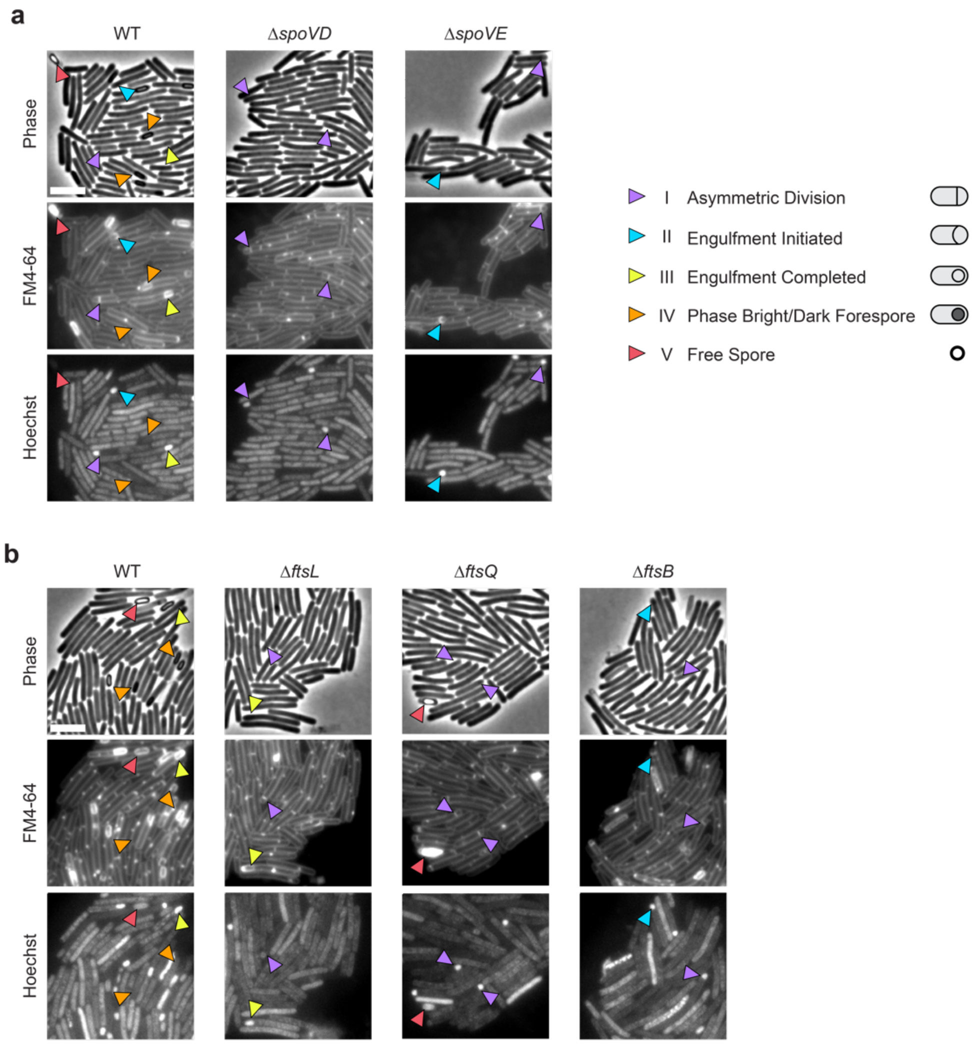


**Supplementary Fig. 5 | Cytological profiling of sporulating cells. a, b,** Representative phase-contrast and fluorescence micrographs of cells sampled from sporulation-inducing 70:30 plates after 18-20 (a) or 20-22 (b) hours of growth. Scale bars, 5 μm. For quantified data, see Fig. 2 c,d and Fig. 3 c,d corresponding to a, and b, respectively. The nucleoid was stained using Hoechst, and the cell membrane was stained using FM4-64. Cytological profiling and scoring of individual cells according to five distinct stages of sporulation are indicated. Purple arrows denote cells undergoing asymmetric division (Stage I) as indicated by a flat polar septum; blue arrows denote cells undergoing engulfment (Stage II) indicated by a curved polar septum; yellow arrows denote cells that have completed engulfment (Stage III) indicated by bright-membrane staining around a fully engulfed forespore; orange arrows denote cells with forespores completing maturation (Stage IV) observable as phase-dark or phase-bright forespores associated with the mother cell in phase-contrast; red arrows denote phase-bright mature free spores (Stage V) observable in phase-contrast. Note that the bright focus in forespores visible with Hoechst DNA staining indicates concentrated nucleoid pumped into the forespore during and after asymmetric division. The membrane and DNA of forespores become inaccessible to staining after completion of engulfment and prior to free spore maturation (Pereira et al., 2013).

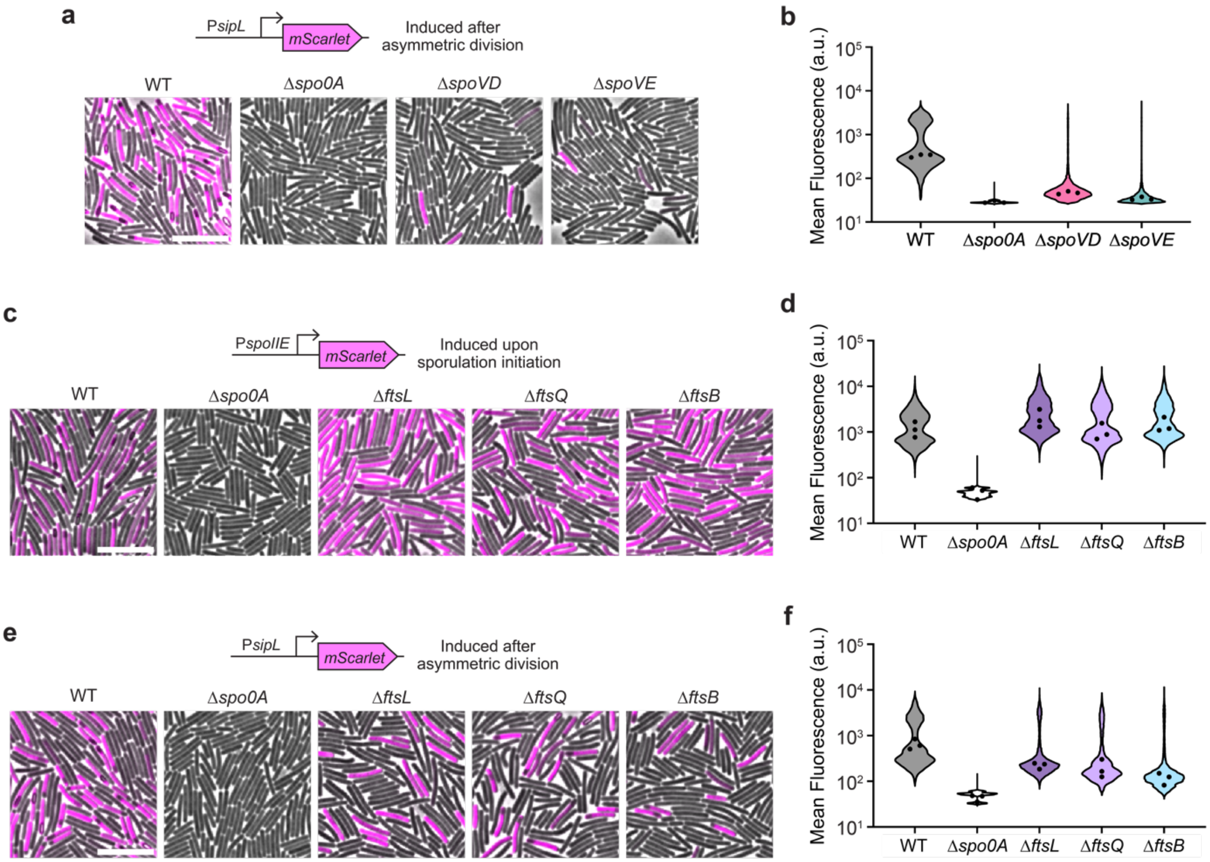


**Supplementary Fig. 6 | Differential activation of downstream sporulation-specific sigma factors in mutants with defects in asymmetric division. a, c, e,** Representative merged phase-contrast and fluorescence micrographs visualizing $P_{spoIIE}$::*mScarlet* (c) and $P_{sipL}$::*mScarlet* (a, e) transcriptional reporters in sporulating cells. $P_{spoIIE}$ is induced immediately upon sporulation initiation, and $P_{sipL}$ is induced after the completion of asymmetric division (Fimlaid et al., 2013). The Δ*spo0A* strain was used as a negative control because it does not initiate sporulation. Scale bars, 10 µm. **b, d, f,** Corresponding violin plots showing quantified mean fluorescence intensities. Black dots represent median values from each replicate. Data from three independent experiments; >3,000 cells per sample.

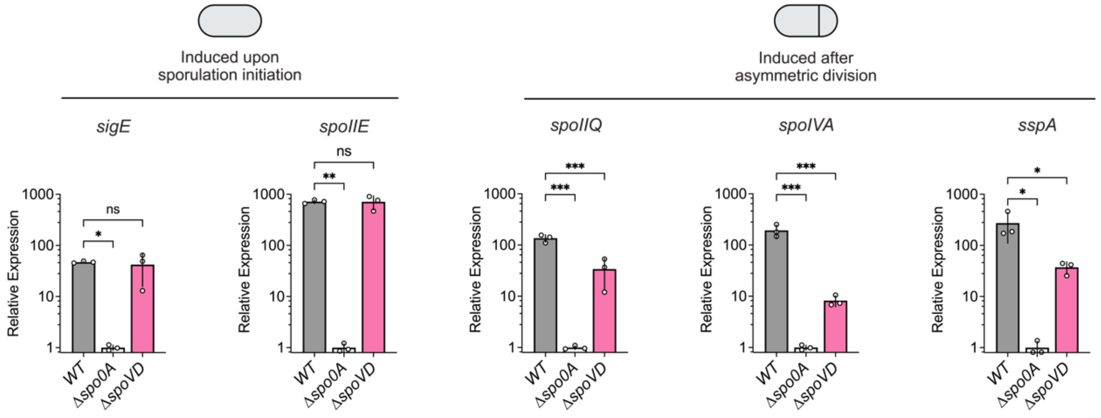


**Supplementary Fig. 7 | Gene expression analyses of sporulation genes in the *spoVD* mutant.** Comparison of representative sporulation gene expression as determined by RT–qPCR. WT, Δ*spo0A,* and Δ*spoVD* cells were sampled from sporulation-inducing 70:30 plates after 10-11 hours of growth. The Δ*spo0A* strain was used as a negative control because it does not initiate sporulation. Transcript levels were calculated relative to the housekeeping gene *rpoB* and normalized to the Δ*spo0A* mean. White dots represent data from each replicate, bars indicate the mean, and error bars indicate standard deviation; data from 3 biological replicates. ns = not significant, *p<0.05, **p<0.01, ***p<0.001; statistical significance was determined using an ordinary one-way ANOVA with Dunnett's test.

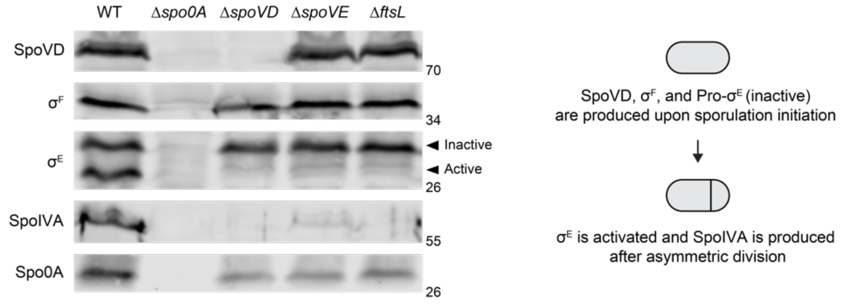


**Supplementary Fig. 8 | Protein level analyses of sporulation proteins in the *spoVD*, *spoVE*, and *ftsL* mutant.** Western blot comparing the levels of the sporulation proteins SpoVD, $\sigma^F$ (SigF), $\sigma^E$ (SigE), SpoIVA, and Spo0A in WT, Δ*spo0A*, Δ*spoVD*, Δ*spoVE*, and Δ*ftsL* cells sampled from sporulation-inducing 70:30 plates after 17 hours of growth. The Δ*spo0A* strain was used as a negative control because it does not initiate sporulation. Spo0A levels reflect the level of sporulation induction (Fimlaid et al., 2013). SpoVD, $\sigma^F$, and $\sigma^E$ are produced upon Spo0A activation at the onset of sporulation. Proteolytic activation of $\sigma^E$ occurs in the mother cell compartment after the completion of asymmetric division. SpoIVA production is dependent on $\sigma^E$ activation. Similar results were obtained from multiple experiments.

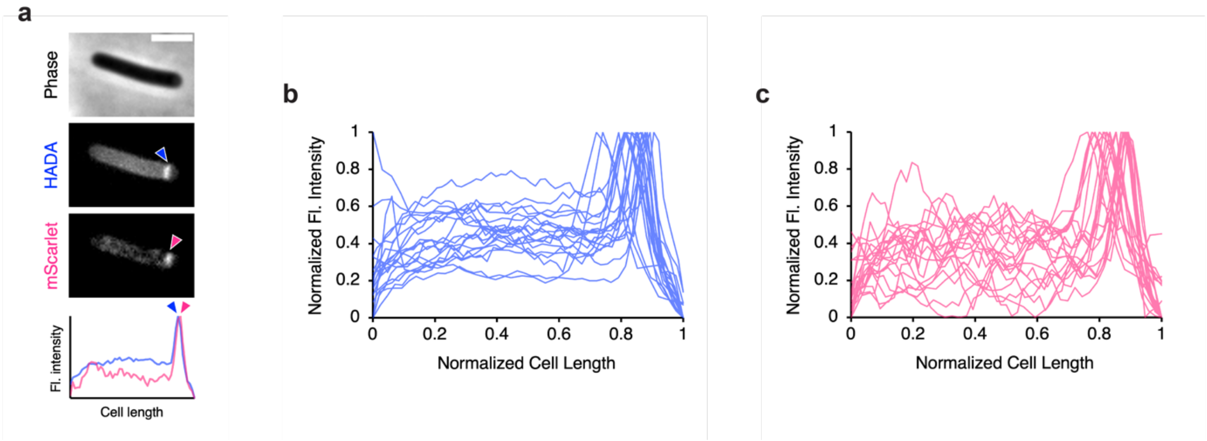


**Supplementary Fig. 9 | SpoVD localizes to the polar septum during asymmetric division. a,** Representative phase-contrast and fluorescence micrographs showing PG incorporation and subcellular localization of mScarlet-SpoVD in a single cell undergoing asymmetric division. Scale bar, 2.5 µm. PG was labeled by incubation with HADA, and *mScarlet-spoVD* was expressed using the native *spoVD* promoter from an ectopic locus. The line graph (bottom) shows the normalized fluorescence (Fl.) intensity of HADA (blue) and mScarlet-SpoVD (pink) along the normalized length. Peaks (arrows) correspond to polar septum sites during asymmetric division. **b, c,** Line graphs showing the normalized fluorescence (Fl.) intensity of the HADA (blue, b) and mScarlet-SpoVD (pink, c) signal along the normalized length of 20 asymmetrically dividing cells sampled from a sporulation-inducing 70:30 plate after 14 hours of growth.

| Organism | Protein | Gene | Essentiality | Domain Schematic | Identity | Similarity | Gaps |
|---|---|---|---|---|---|---|---|
| *C. difficile* str. 630 | FtsL | *cd630_26570* (*ftsL*) | Non-essential | | 100% | 100% | 0% |
| *B. subtilis* str. 168 | FtsL | *ftsL* | Essential | | 22% | 43% | 27% |
| *E. coli* str. K-12 | FtsL | *ftsL* | Essential | | 12% | 32% | 35% |
| *C. difficile* str. 630 | FtsQ | *cd630_26500* (*ftsQ*) | Non-essential | | 100% | 100% | 0% |
| *B. subtilis* str. 168 | DivIB | *divIB* | Essential | | 21% | 38% | 32% |
| *E. coli* str. K-12 | FtsQ | *ftsQ* | Essential | | 15% | 33% | 34% |
| *C. difficile* str. 630 | FtsB | *cd630_34920* (*ftsB*) | Non-essential | | 100% | 100% | 0% |
| *B. subtilis* str. 168 | DivIC | *divIC* | Essential | | 18% | 35% | 35% |
| *E. coli* str. K-12 | FtsB | *ftsB* | Essential | | 22% | 41% | 16% |

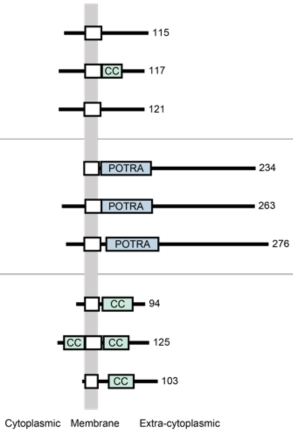


**Supplementary Fig. 10 | Comparison between *C. difficile*, *B. subtilis*, and *E. coli* orthologs of divisome-associated FtsL, FtsQ/DivIB, and FtsB/DivIC proteins.** Predicted domains were identified using InterPro (Paysan-Lafosse et al., 2023). Domain schematics are scaled according to amino acid lengths. White boxes represent transmembrane domains or signal peptides; green boxes represent coiled-coil domains (CC); blue boxes denote polypeptide-transport-associated domains (POTRA) found in FtsQ/DivIB proteins. Pairwise protein sequence alignment data compared to the *C. difficile* ortholog of each protein were calculated using EMBOSS Needle (Madeira et al., 2022). Despite low sequence conservation among FtsQ/DivIB and FtsB/DivIC orthologs, they are expected to be functionally equivalent and contain similar domain compositions. The distinction between the Div and Fts nomenclature was previously suggested to distinguish between orthologs found in Gram-positive and Gram-negative bacteria, respectively. However, the Div nomenclature has only been adopted for members of Bacilli, while non-Firmicutes Gram-positive bacteria such as members of Actinobacteria use the Fts nomenclature. Furthermore, the genomic contexts of *ftsL* and *ftsQ*/*divIB* within the *dcw* cluster is widely conserved among Gram-positive (e.g., Bacilli, Clostridia) and Gram-negative (e.g., Negativicutes) members of Firmicutes (Fig. 1b) as well as other bacterial phyla (Garcia et al., 2021; Megrian et al., 2022). These observations highlight the arbitrary nature of the distinction between Fts and Div proteins. Hence, we suggest the adoption of Fts nomenclature for all bacteria, including members of Bacilli, for clarity and consistency. Accordingly, we have applied the Fts nomenclature to the *C. difficile* orthologs in this study.

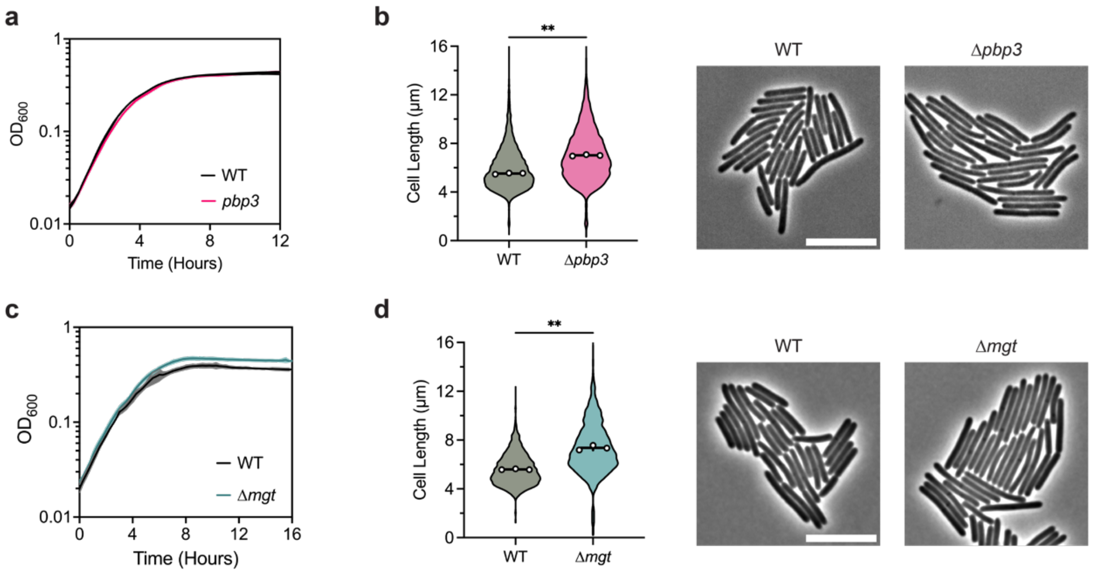


**Supplementary Fig. 11 | Non-essential PG synthases PBP3 and MGT are dispensable for growth and division. a, c,** Growth profile of Δ*pbp3* (a) and Δ*mgt* (c) compared to WT. Data from a single growth curve experiment; mean, and standard deviation are plotted from three biological replicates. **b, d,** Representative phase-contrast micrographs (right) and violin plots (left) showing cell length distributions of WT, Δ*pbp3* (b), and Δ*mgt* (d) cells sampled from BHIS cultures during exponential growth ($OD_{600}$ ~0.5). White circles indicate means from each replicate, black line indicates the average mean. Data from three biological replicates; >3,000 (b) and >500 (d) cells per sample. **$p<0.01$; statistical significance was determined using a paired *t* test. Scale bars, 10 µm.

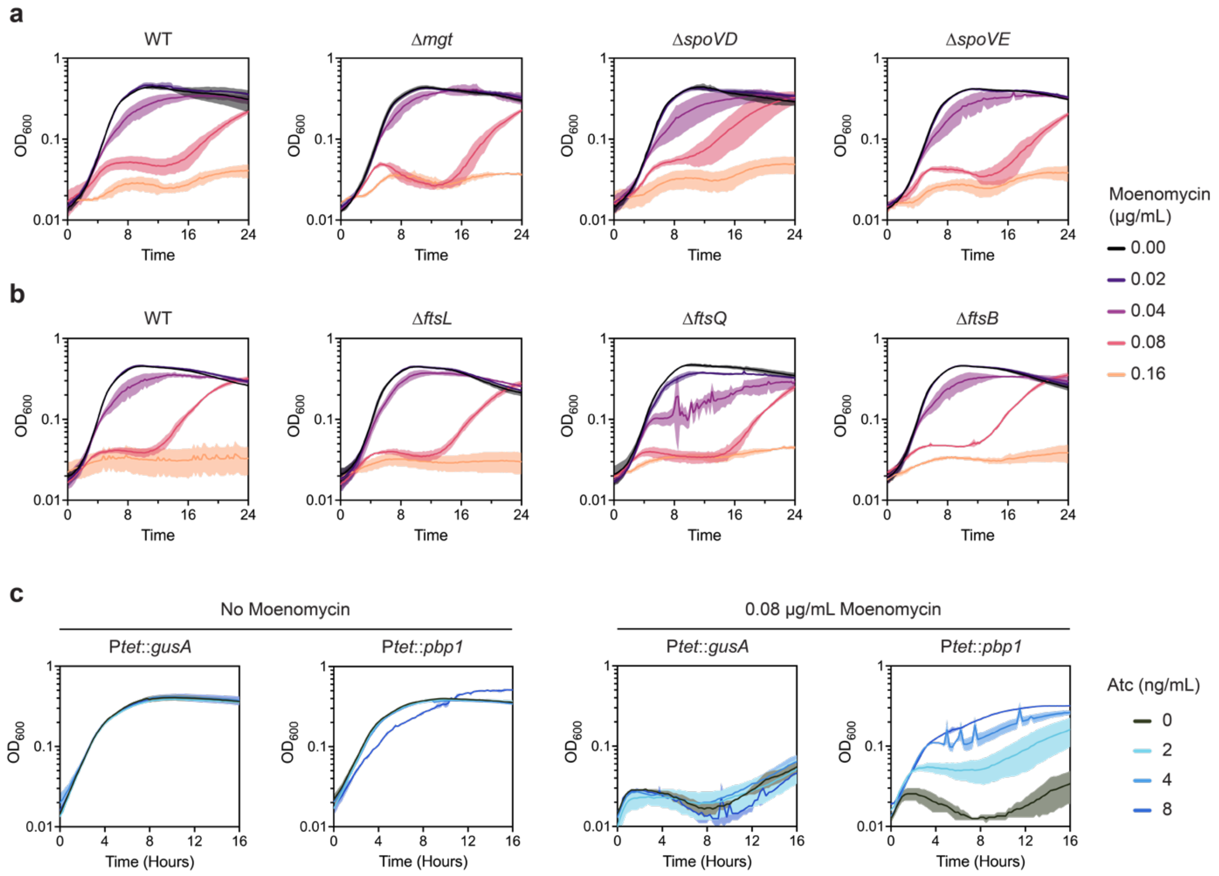


**Supplementary Fig. 12 | Sensitivity to moenomycin is counteracted by *pbp1* overexpression but unaffected by the absence of non-essential PG synthases and divisome components. a, b,** Growth profiles of WT and various mutants in BHIS supplemented with different concentrations of moenomycin as indicated. Data from a single growth curve experiment; mean, and standard deviation are plotted from three biological replicates. **c,** Growth profiles of WT strains harboring a *pbp1* overexpression (P*tet*::*pbp1*) or negative control (P*tet*::*gusA*) vector in BHIS with or without moenomycin. Expression from the anhydrotetracycline (ATc)-inducible promoter (P*tet*) was induced with various concentrations of ATc, as indicated. Data from a single growth curve experiment; mean, and range are plotted from two biological replicates.
